# Not just flowering time: A resurrection approach shows floral attraction traits are changing over time

**DOI:** 10.1101/2022.08.18.504321

**Authors:** Sasha G.D. Bishop, Shu-Mei Chang, Regina S. Baucom

## Abstract

Contemporary anthropogenic changes in climate and landscape form a complex set of selective pressures acting on natural systems, yet, in many systems, we lack information about both whether and how organisms may adapt to these changes. In plants, research has focused on climate-induced changes in phenology and the resultant potential for disruption of plant-pollinator interactions, however there remains a paucity of knowledge regarding how other pollinator-mediated traits may be involved in adaptive response. Here, we use resurrection experiments to investigate the phenotypic basis of adaptation in a mixed-mating system plant, the common morning glory (*Ipomoea purpurea*). Specifically, we measure temporal and spatial changes in traits grouped into three categories relevant to plant-pollinator interactions - floral morphology, floral rewards, and floral phenology. We show a significant temporal increase in corolla size and shift to earlier flowering times, as well as a potential for increased investment in floral rewards, all of which are driven primarily by populations at more northern latitudes. Additionally, we find evidence for directional selection on floral morphology and phenology, and evidence of balancing selection acting on anther-stigma distance. Overall, these results show an adaptive response in line with greater investment in pollinator attraction rather than self-pollination and fine-scale spatial differences in adaptive potential.

## INTRODUCTION

Climate change is causing rapid, simultaneous changes in several environmental variables, consequently exposing communities to novel combinations of abiotic and biotic conditions. These changes include altered temperature, precipitation, photoperiod, CO_2_ and N_2_O levels, many of which display seasonally and geographically distinct patterns, and all of which together form a complex and multifactorial suite of selective pressures (IPCC 2022). For most species, we have very little understanding of which traits may underlie adaptive responses when exposed to the multivariate selective pressures typical of climate change (Abatzoglou et al. 2020; Gallagher and Campbell 2021). In plant systems, the potential that climate change may disrupt plant-pollinator interactions is of particular concern. This is because many insect pollinators have faced significant global declines (Potts *et al*., 2010; Winfree *et al*., 2011; Thomann *et al*., 2013; Hallmann *et al*., 2017; Soroye *et al*., 2020), and these declines have been accompanied by concomitant reductions in insect-pollinated plants (Biesmeijer *et al*., 2006).

Much of the research investigating climate-induced disruptions of plant-pollinator interactions has focused on flowering phenology, with studies typically showing a general trend of earlier flowering across species (Byers, 2017; Renner & Zohner, 2018; Gérard *et al*., 2020; Soares *et al*., 2021). However, plant-pollinator interactions are not mediated by phenology alone, but by an array of interacting traits influencing both rate of pollinator visitation and pollinator effectiveness (Glenny *et al*., 2018; Sletvold, 2019). For example, pollinator preference positively correlates with corolla size (Galen, 2000; Chapurlat *et al*., 2020), and plant-pollinator interactions are often mediated by floral rewards received by pollinators in the form of nectar and pollen (Eckert *et al*., 2010; Campbell & Powers, 2015; Descamps *et al*., 2018, 2021; Parachnowitsch *et al*., 2018). Although standing genetic variation of these floral traits is frequently high in field settings, demonstrating a potential for rapid evolutionary shifts (Thomann *et al*., 2015), strikingly few studies have investigated changes in suites of pollinator-mediated traits beyond flowering phenology, such as corolla size and/or traits associated with floral rewards.

Traits related to self-pollination can also evolve given climate shifts (Van Etten & Brunet, 2013), with some evidence pointing to increased selfing as climate change and/or pollinator declines associated with land use changes (Eckert *et al*., 2010; Jones *et al*., 2013; Cheptou, 2019). A generalized expectation for hermaphroditic plant species in this regard is a shift to smaller anther-stigma distances – *i*.*e*., decreased distance between the anthers and stigmas within perfect flowers (Chang & Rausher, 1998) – since smaller anther-stigma distance is highly correlated with greater self-pollination (Chang & Rausher, 1998), as well as decreased investment in pollinator attraction traits such as flower size and nectar quality (Levin 2010). However, there is a major gap in our understanding of how traits that are crucial for plant-pollinator interactions may be evolving over time as a response to a changing climate, and a number of predictions could be made. Are traits responsible for plant-pollinator interactions evolving in light of pollinator decline, such that plants are evolving greater floral displays to attract pollinators? Or are traits that promote selfing like lower anther-stigma distance evolving to maintain populations in light of reduced pollinator presence?

In this work, we compare floral traits of populations of *Ipomoea purpurea* (common morning glory) stored as seed for a number of years to that of contemporary populations (*i*.*e*. a resurrection approach) to examine the potential that traits responsible for plant-pollinator interactions and self-fertilization may be evolving over time and in light of global change. Specifically, we used three separate common garden greenhouse studies to compare floral traits of populations sampled in 2003 to those of the same populations sampled nine years later in 2012. Populations were located across a large range of the southeast and midwest US, such that we examined floral morphology and flowering phenology trends across both different collection times and spatial locations. We measured traits grouped into three classes relevant to plant-pollinator interactions – floral morphology, floral phenology, floral rewards – and addressed the following questions: (1) Is there evidence of variation in floral traits and do those traits exhibit any changes between sampling years (2003 vs. 2012)? and (2) Are changes in phenotype likely the result of adaptation through natural selection or neutral processes? Our expectation is that an adaptive response toward greater selfing will result in selection for decreased anther-stigma distance and smaller flower size, whereas positive changes in floral size and rewards indicate greater investment in pollinator attraction. Cumulatively, our results fill critical gaps in our understanding of plant responses to environmental change by highlighting adaptive changes in floral traits beyond phenology and providing evidence of small-scale spatial heterogeneity in adaptive potential.

## MATERIALS AND METHODS

### Study System & Sampling History

*Ipomoea purpurea* (Convolvulaceae), or the common morning glory, is an annual, weedy vine widely distributed across the eastern, midwestern, and southern United States (Tiffin & Rausher, 1999). It is frequently found along roadsides or in agricultural settings, often in areas of high disturbance (Tiffin & Rausher, 1999). The species employs a mixed mating system as it outcrosses ∼50% of the time (Kuester *et al*., 2017) and is typically pollinated by bees, syrphid flies, and wasps. *Ipomoea purpurea* germinates in late spring and typically begins flowering after 6-8 weeks of growth. Flowering continues until the first frost, and the fruits are dehiscent capsules that contain between one to six seeds.

In this resurrection study, we used replicate seeds sampled at two time points (2003 and 2012) from 23 different populations located within agricultural fields from Tennessee and North and South Carolina in the US. Details of the sampling are presented in (Kuester et al. 2016). Briefly, at each sampling time point, a 10-30 m transect was drawn and seeds from a single plant were collected at 1-2 m intervals down the transect. Agricultural crops within sampled fields altered between soy and maize between 2003 and 2012 as is typical of crop rotation schemes, however GIS images indicated that no major changes in land-use occurred between sampling years. Thus, the collections from these populations represent time series data that capture environmental and phenotypic changes from the combination of climatic or agricultural regime changes.

### Greenhouse Experiments

#### Floral Morphology

To assess floral morphology, we planted replicates of field-collected seeds from maternal lines sampled from 15 populations (Fig. 1; STable 1) distributed from the Cumberland Plateau of central Tennessee to the Coastal Plain region of North and South Carolina and from two collection times (2003 and 2012). Specifically, we planted seeds from 6-18 maternal lines per population for 2003 (average = 14.67, median = 16) and 1-29 maternal lines per population for 2012 (average = 15.8; median = 15; see Table S1 for the exact number of maternal lines per population). Seeds were scarified and planted in 4-inch pots which were arranged in a completely randomized design at the Matthaei Botanical Gardens (MBGNA) at the University of Michigan (Ann Arbor, MI, USA). This experiment was performed in 2015.

**Figure 1.**
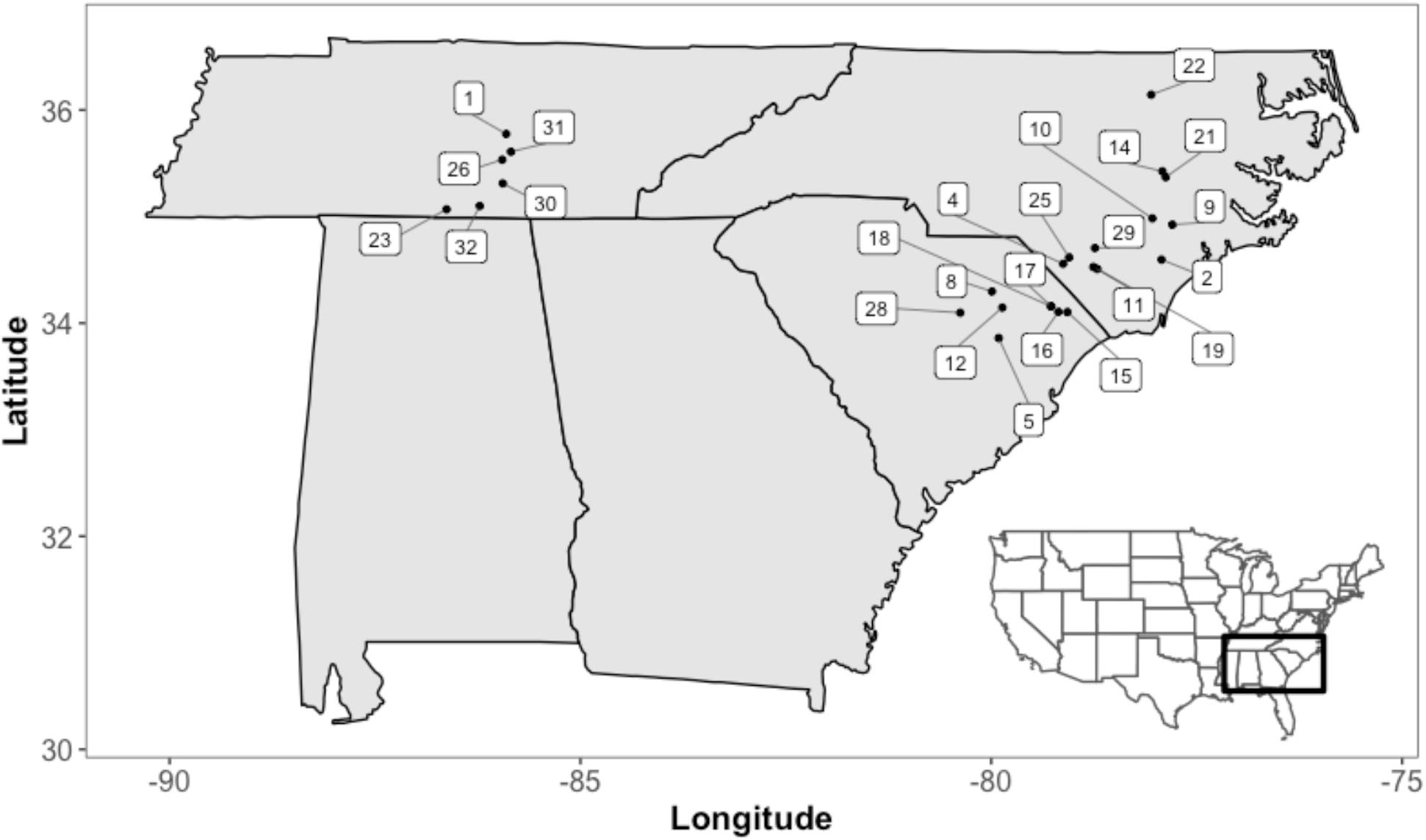
Distribution of sampling localities of *I. purpurea* labeled with population number. All populations were sampled from the edge of agricultural soy and maize fields. Fifteen populations were included in the resurrection experiment looking at floral morphology, twenty-three populations for phenology, and four populations with in-depth maternal line sampling were used to measure floral reward traits. Population representation for each resurrection experiment can be found in Supplemental Table 1.

Floral traits (corolla width (cm), corolla length (cm), and height of the tallest anther (cm) and the pistil (cm) for an estimate of the anther-stigma distance, ASD) on an average of 6 flowers/plant were measured using digital calipers. Measurements were spread across 17 sampling dates with an average of 2.3 flowers measured per plant on each date, such that flowers from the 2003 and 2012 cohorts were always measured at the same time, ensuring equi-aged flowers. In total, 2836 flowers were measured from 456 plants. Corolla width was measured as the diameter of a fully open corolla, corolla length as the distance from the rim of the corolla to where it fuses with the receptacle, and anther-stigma distance (ASD) as the difference between the height of the pistil and the tallest stamen.

#### Floral Phenology

To assess floral phenology, we performed a separate common garden experiment in 2013 at the University of Georgia Plant Biology Greenhouses (Athens, GA, USA) using field collected seeds from 23 populations again from two different years (2003 and 2012). A total of 451 plants were included in this study, with 2-16 plants per population (mean = 9.8, median = 11). Thirteen of these populations overlapped with those included in the greenhouse experiment at MBGNA assessing floral morphology (Fig. 1). We recorded the first occurrence of a fully open bloom as the date of first flower for all experimental individuals. To determine if there were size differences between plants from different sampling years, we counted the number of leaves of each individual, and dried plants at 70°C for three days and weighed each individual for an estimate of dried biomass. Germination of seeds in this experiment ranged from 50-98% across populations and varied between years, with more seeds germinating from the 2003 collections compared to the 2012 collections (87% vs 84%, p < 0.001; Kuester et al. 2016).

#### Floral Rewards

To measure floral rewards, we replanted a subset of four populations (Supp. Table 1) in a separate experiment at MBGNA in 2017 to quantify the number of pollen grains produced and the nectar sucrose content (°Brix), which we consider an important component of total nectar reward. We planted replicates of eight maternal lines for each of these populations, again sampled both in 2003 and 2012 (except for Duplin East from 2012 which included only 6 maternal lines). We measured a total of 1468 flowers from 213 plants, with an average of 26.6 plants per population and 6.89 flowers per plant.

To extract nectar from the flower, 10uL of reverse osmosis (RO) water was pipetted directly into the base of a flower, pushing the pipette tip past the base of the stamens and pipetting up and down to mix and extract nectar. We then quantified sucrose content of this nectar/water solution using a pocket refractometer to record percent mass sucrose (°Brix, hereafter nectar sucrose content). We counted pollen by removing the second tallest anther in each flower with forceps near the time of anthesis (i.e. early morning) when pollen was mature. We then gently brushed the anther against all four corners of a basic fuchsin gelatin cube (Beattie, 1971). The cube was placed on a glass microscope slide, heated on a 180°C hot plate until the cube completely melted, covered with a cover slip, and imaged with an iPhone camera affixed to a light microscope. We obtained a pollen count by analyzing pollen slides using the Analyze Particles function in ImageJ (Schneider *et al*., 2012) with the default particle size setting (0-150).

### Data Analysis

#### Temporal and Spatial Effects on Floral Traits

We first examined possible phenotypic evolution by comparing differences in trait distributions between collection years using a Kolmogorov-Smirnov test across all populations. To determine if the mean trait values were different between sampling years and spatial locations, we performed a linear mixed model using the lme4 package v. 1.1.29 (Bates *et al*., 2015) in R (v. 4.2.0; R Core Team 2022) with year, latitude, and the interaction of year and latitude as fixed effects and population identifier as a random effect to control for longitudinal differences. Each phenotypic trait was used as the dependent variable in separate models of the following general form:

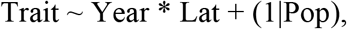

and we used the bestNormalize v. 1.8.2 package (Peterson 2021) to determine the appropriate transformation for each trait to adhere to assumptions of normality. We assessed the significance of effects using the anova() function from lmerTest v. 3.1.3 (Kuznetsova *et al*., 2017), which performs a type III ANOVA and uses the Satterthwaite method to determine the degrees of freedom. The day of first flowering showed a bimodal distribution (see Results); while most experimental individuals flowered in the first wave, a small group of individuals flowered for the first time in what we describe as a second wave. Due to the resulting bimodal distribution of first flowering dates (Fig. 1), a normality transformation was not appropriate. We thus elected to model each flowering wave separately. Moving forward, we focus our statistical analysis primarily on the first wave of flowering, as that captures information about flowering phenology for the majority of individuals in the experiment. However, we do describe differences between the first and second waves of flowering in the discussion for illustrative purposes.

The floral reward traits (pollen number and Brix) were measured on replicate maternal lines from four populations. Thus, we included a maternal line effect in our ANOVAs when testing for temporal changes in floral reward traits, and we used least square means to assess the potential for temporal changes within each population separately. For both traits, maternal line and population were included as random effects with year, latitude, and the interaction between year and latitude included as fixed effects:

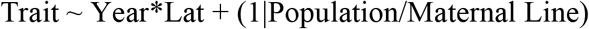

Data were again transformed to meet assumptions of normality and analyzed with a type III ANOVA.

#### Phenotypic Evolution

We next performed a screen to determine if the dominant evolutionary force influencing floral traits over time was selection, migration, or drift. To do so, we compared the change in trait value (*𝛿t*; or 2012 – 2003) to the initial value in 2003 (*t*), and assessed potential changes in the variance of each trait, following (Goldberg *et al*., 2020). Our expectations are presented in Table 1, but, briefly, the presence of selection influencing trait change would be evident by the following: a significant change in mean trait value from 2003 to 2012, a reduction in trait variation, and significant non-zero slope between *t* and *𝛿t. 𝛿t* values either above or below zero would support positive or negative directional selection, respectively. If drift were driving phenotypic evolution, we would expect both an increase in trait variation across populations and a zero-slope relationship between *𝛿t/t*. No net change in trait value or variance would suggest the presence of balancing or disruptive selection in traits; if values for *𝛿t* were scattered both above and below zero, a significant linear regression between *t* and *𝛿t* with a positive slope that intercepts with the line *𝛿t*=0 would suggest disruptive selection (i.e. small *t* has a negative *𝛿t*; large *t* has a positive *𝛿t*), while a negative slope would indicate balancing selection (i.e. small *t* has a positive *𝛿t*; large *t* has a negative *𝛿t*). Finally, balancing selection can be differentiated from the homogenizing force of migration since migration would be expected to decrease trait variation.

**Table 1.** A framework adapted from (Goldberg *et al*., 2020) to differentiate between drift, migration, and selection on trait evolution between sampling years. *t* refers to the least squares mean for a trait value, 𝛿*t* is the difference in mean trait value from 2012-2003, and the PCV is used to assess variance in *t*. In addition to the expectations outlined by Goldberg *et al*., 2020, for balancing and disruptive selection, we expect 𝛿*t* values to be distributed around 0 such that the direction of change (increase or decrease) is dependent on the starting value, whereas for directional selection, the direction of change in 𝛿*t* will remain consistent regardless of starting value.

| | $\delta t$ | $\delta t$ versus $t$ | Variance in $t$ |
| --- | --- | --- | --- |
| **Drift** | No net change | Slope = 0 | Increase |
| **Migration** | No net change | median $\delta t \approx 0$ | Decrease |
| **Directional Selection (+)** | Positive | $\delta t > 0$, positive slope | Decrease |
| **Balancing Selection** | No net change | median $\delta t \approx 0$, negative slope | No change |
| **Disruptive Selection** | No net change | median $\delta t \approx 0$, positive slope | No change |

To apply the (Goldberg *et al*., 2020) framework to our system, we calculated trait variation as the phenotypic coefficient of variation (PCV; (standard deviation(*x*)/mean(*x*)) 100%; where *x* is the trait of interest). To test for temporal changes in PCV, we used the Coefficient of Variance with Confidence Intervals (cvcqv) package v. 1.0.0 in R (Beigy, 2019) and used bootstrap resampling to obtain confidence intervals, then conducted a two-sided independent t-test for each trait. We used a linear regression assessed with a type II ANOVA and Pearson’s correlation coefficient to determine if variation in the change in trait over time (*𝛿t*) was explained by the initial trait value (*t*).

Finally, we revisited evidence for selection based on the relationship between *t* and *𝛿t* by including latitude as a potential predictor of *𝛿t*. The predictions for *𝛿t/t* outlined above focus on detection of evolutionary forces that are consistent across populations, resulting in an overall dominant effect on the species under consideration. However, climatic changes can vary dramatically across latitude, resulting in different selective forces over space. Based on preliminary analysis, changes in corolla width appeared stronger in northern latitudes, with a significant latitude year interaction when assessing mean changes in this trait. Thus, we also included a latitude effect for traits when examining the relationship between *𝛿t/t*.

## RESULTS

### Temporal and Spatial Effects

Patterns of trait change between collection years varied across floral morphology, phenology, and floral reward traits. The trait distribution for corolla width was significantly different between collection years (two-sample D = 0.157, p = 1.04e-14, Fig. 2), and this difference was reflected in a change in mean value, with corollas becoming significantly wider over time (4.5 cm in 2003 vs 4.8 cm in 2012; F = 7.093, numDF = 1, denDF = 12.10, p = 0.020; Table S2). Although it appeared that corolla width increased across most populations (Figure 3), we found a highly significant interaction between latitude and year (F = 23.388, numDF = 1, denDF = 519.82, p = 1.75 × 10^−6^; Table S2) and a highly significant effect of latitude (F = 16.850, numDF = 1, denDF = 2662.85, p = 4.167 × 10^−5^; Table S2) such that the change in corolla width was greater in populations at more northern latitudes (STable 2, Figure 3). No change from 2003 to 2012 was detected in plant biomass (t = 0.078, df = 54.289, p = 0.938) or in pre-flowering leaf count (t = 0.1865, df = 704.31, p-value = 0.8521), suggesting that increased corolla width detected here is not due to an overall increase in plant size.

**Figure 2.**
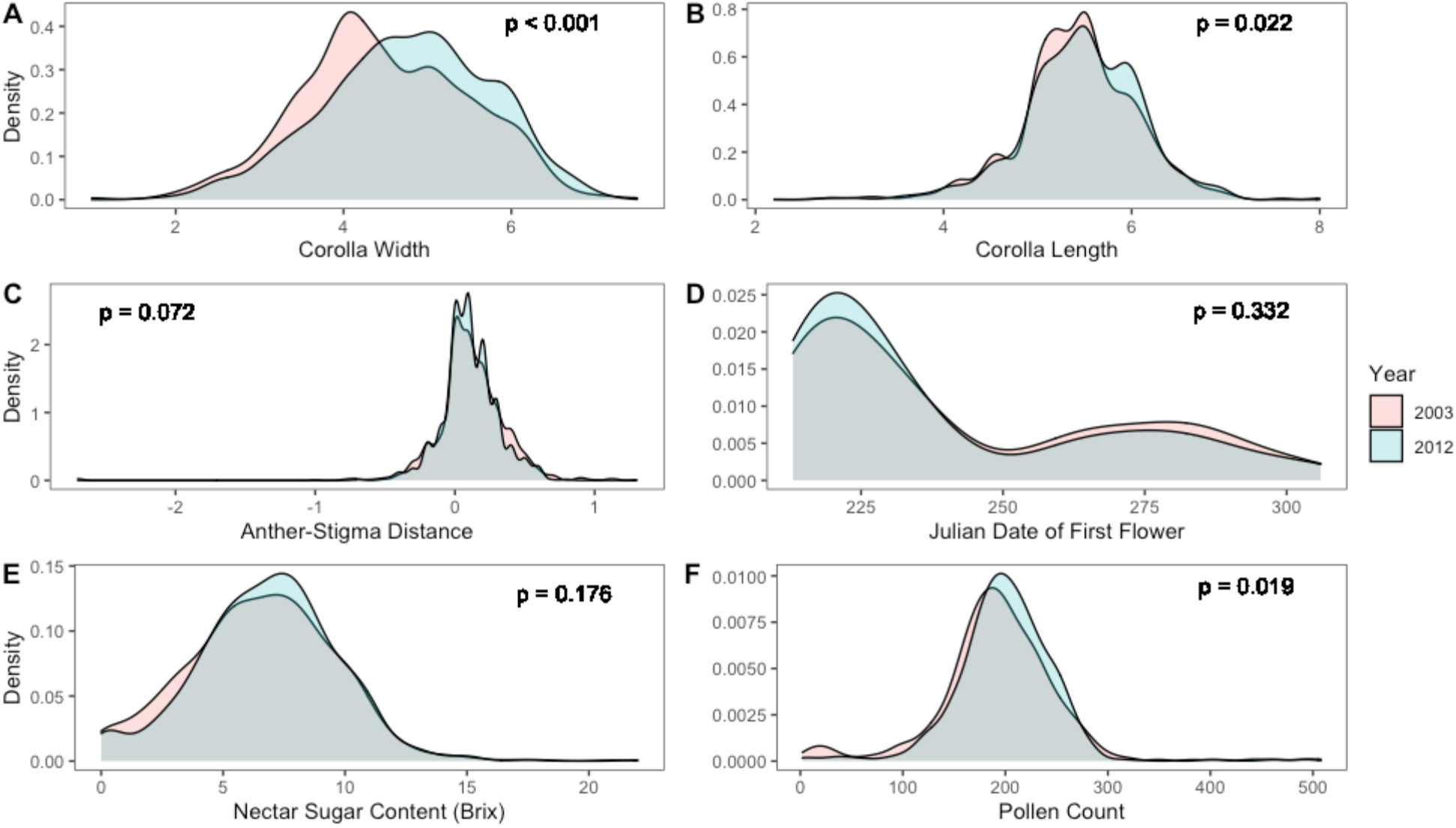
Distribution of trait values across all populations in 2003 and 2012. P-values come from Kolmogorov-Smirnov tests for each trait.

**Figure 3.**
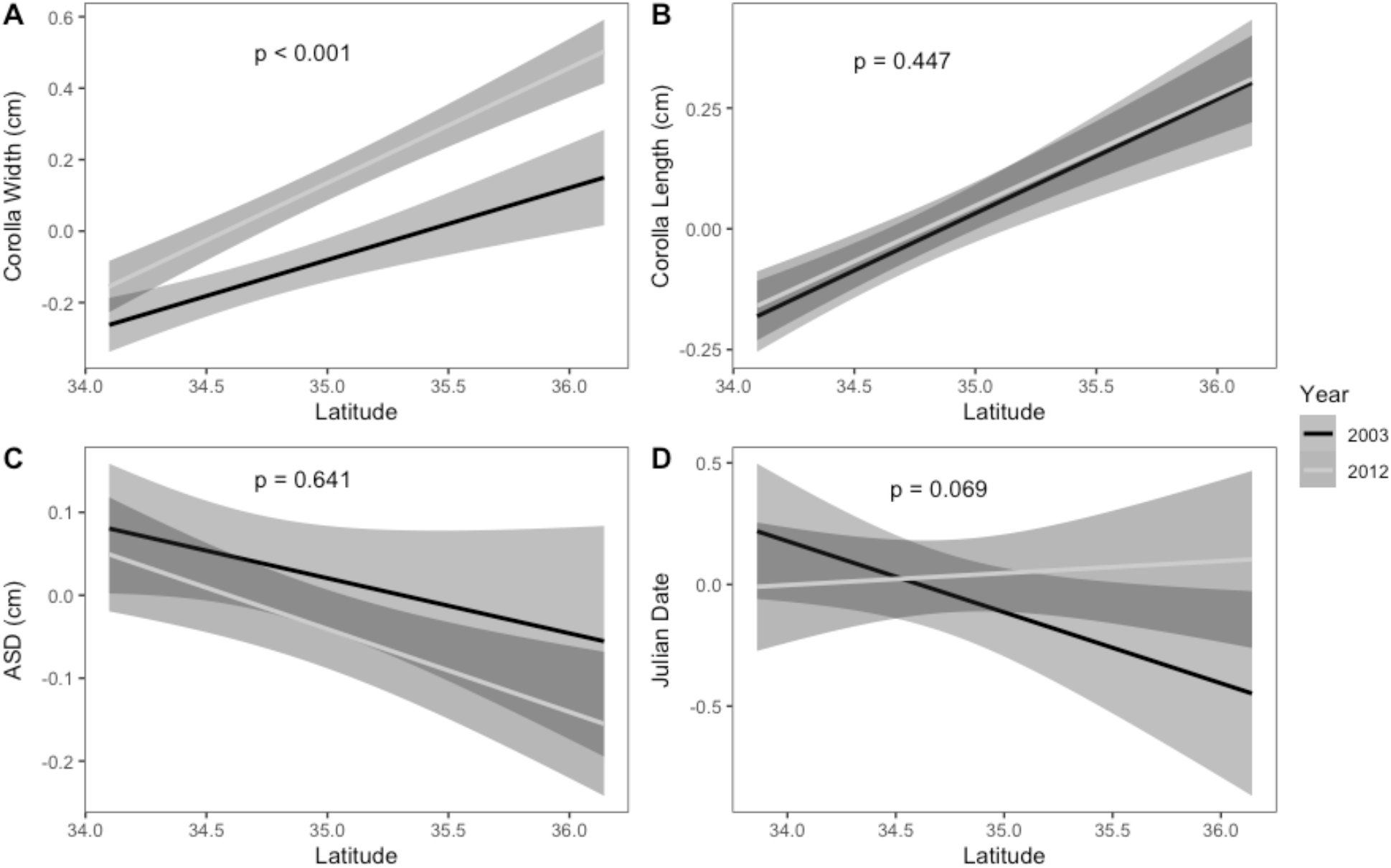
Linear mixed models for population means across latitude for a) corolla width, b) corolla length, c) anther-stigma distance, and d) the Julian date of first flower for the first wave of flowering. Each line is plotted with a 95% confidence interval and p-values on plots refer to the year*latitude effect from a type III ANOVA of linear mixed model 1.

Compared to corolla width, a smaller shift occurred in the distribution of corolla length (two-sided D = 0.057, p = 0.022; Fig. 2), and we found evidence of a slight but significant increase in corolla length between collection years (5.43 cm in 2003 vs 5.47 cm in 2012; F = 10.472, numDF = 1, denDF = 11.77, p = 0.007; Table S2). There was no indication that corolla length differed across latitudes (F = 0.041, numDF = 1, denDF = 2781.94, p = 0.840; Table S2) nor was there a significant interaction between latitude and year (F = 0.580, numDF = 1, denDF = 923.79, p = 0.447; Table S2). Due to non-normality in the data even after correction, we checked for robustness of the year effect using a permutation test and again uncovered a nearly significant change in corolla length over time (p = 0.069).

For the final floral morphology trait we examined, anther-stigma distance, we found a nearly significant change in trait distribution (two-sided D = 0.049, p = 0.072; Fig. 2), however no evidence for a change in trait mean over time (F = 4.42, numDF = 1, denDF = 9.67, p = 0.659; Table S2), nor did we find evidence of a significant effect of latitude (F = 1.587, numDF = 1, denDF = 2468.51, p = 0.072; Table S2), or significant interaction between year and latitude (F = 2.633, numDF = 1, denDF = 1576.65, p = 0.641; Table S2, Figure 3).

When assessing flowering phenology, we found that the start of flowering occurred in two waves (Fig. 2). For the first wave of flowering onset, we identified an almost significant effect of both collection year (F = 3.300, numDF = 1, denDF = 300.97, p = 0.070; Table S2) and interaction of collection year and latitude (F = 3.339, numDF = 1, denDF = 300.97, p = 0.069; Table S2) on the date of first flower. For the second wave of flowering onset, we found that the day of first flowering of the second wave differed according to latitude (F = 6.194, numDF = 1, denDF = 22. 12, p = 0.021; Table S2), but found no evidence for a collection year effect (F = 0.232, numDF = 1, denDF = 139.97, p = 0.631; Table S2) nor a significant interaction between collection year and latitude for this trait (F = 0.225, numDF = 1, denDF = 139.94, p = 0.636; Table S2, Figure 3).

Like floral morphology and flowering time, we found collection year and latitudinal differences in the floral reward traits. Similar to corolla width, the distribution of pollen grain number exhibited a significant shift in the distribution toward greater pollen grain number in 2012 (two-sided D = 0.100, p = 0.019; Fig. 2). However, we did not find an overall effect on average pollen number between years (F = 0.028, numDF = 1, denDF = 1.90, p = 0.883; Table S2), nor was there a difference according to latitude (F = 2.187, numDF = 1, denDF = 163.41, p = 0.141), or significant interaction between collection year and latitude (F = 2.180, numDF = 1, denDF = 33.25, p = 0.149; Table S2, Figure 4). For nectar sucrose content (°Brix), we found a significant interaction between year and latitude, such that the more northern populations exhibited increased °Brix over time (F = 4.59, numDF = 1, denDF = 60.45, p = 0.036, Table 2; Figure 4). There was no support for an overall year effect for this reward trait (F = 0.003, numDF = 1, denDF = 1.94, p = 0.961) but there was a significant latitude effect (F = 5.877, numDF = 1, denDF = 200.63, p = 0.016).

**Figure 4.**
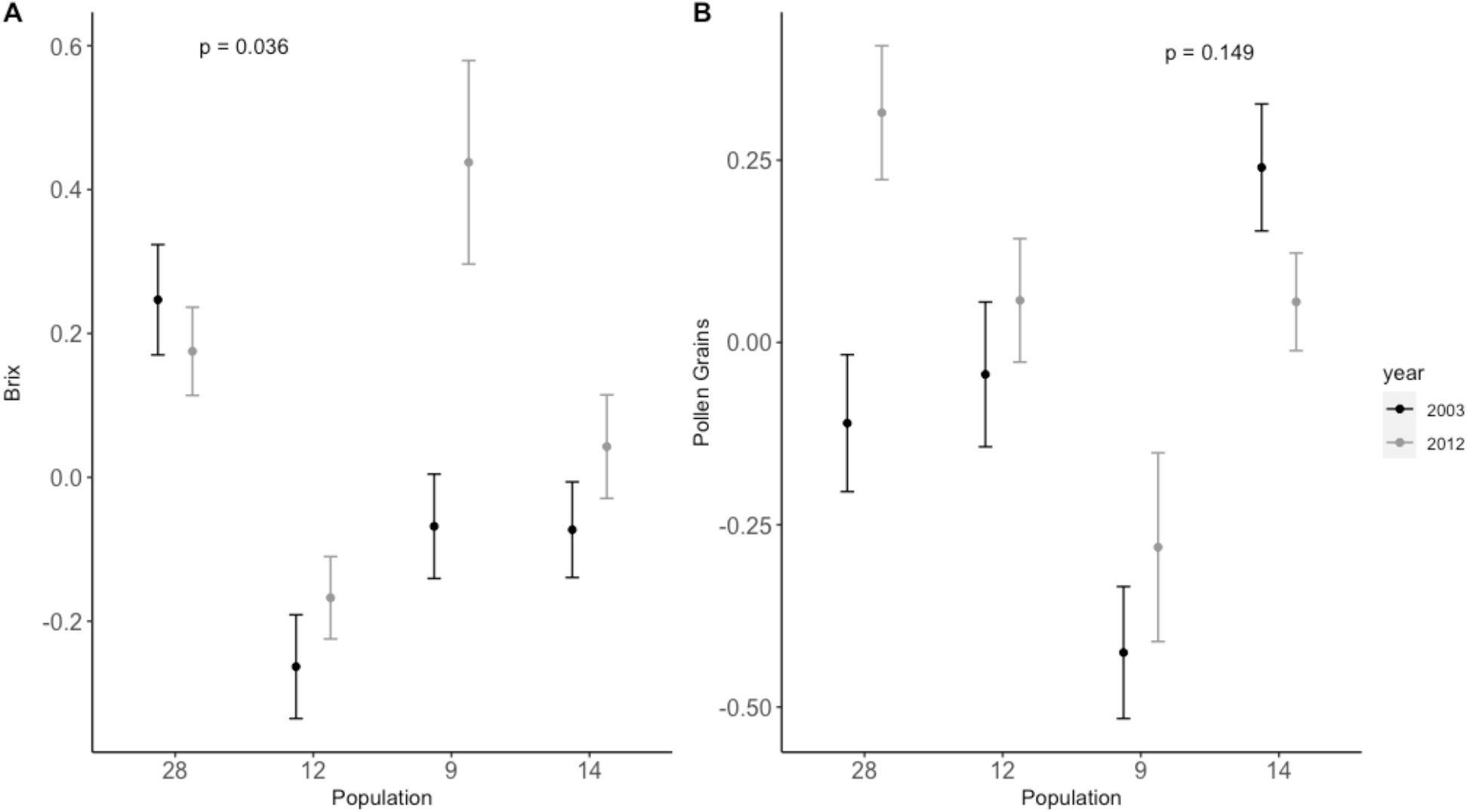
Per population changes in floral rewards from 2003 (black) to 2012 (grey) with populations ordered left to right from lowest to highest latitude. Mean and standard error for °Brix (A) and pollen count (B) are shown for the four measured populations and p-values refer to the year*latitude effect from a type III ANOVA of linear mixed model 1.

### Adaptive Evolution

Using the framework of Goldberg et al 2020 (Table 1), we found that most of the morphology and flowering time traits examined (corolla width, corolla length, anther-stigma distance, and flowering time of the first flowering wave) appeared to be evolving under some form of selection. We did not include pollen number and °Brix in this analysis since four populations is insufficient for a regression analysis.

Corolla width displayed a significant, negatively sloped relationship (R = −0.55, p = 0.04) between the change in mean trait value (*𝛿*t) and starting mean trait value in 2003 (*t*) after the removal of a single outlier population (Figure 5). There was likewise evidence for reduced variation in this trait over time (5.4% reduction in the phenotypic coefficient of variation (PCV), t = 1.854, p = 0.059; Table S3). These two results together, along with the significant increase in trait mean over time, provide mixed evidence for either directional selection (*i*.*e*., reduction in variation and change in mean) or balancing selection (*i*.*e*., relationship between *𝛿*t and t). However, including latitude as an explanatory effect for *𝛿*t in an analysis of variance revealed a significant interaction between latitude and *t* (F = 6.058, numDF = 1, denDF = 11, p = 0.03, Table S2), corroborating previous evidence that latitude plays a strong role in determining changes over time in corolla width. Based on this model, the slope of *𝛿t/t* for corolla width becomes positive above a latitude of 34.9; all populations except one above this latitude also demonstrate *𝛿t* values greater than zero. It thus appears that populations at northern latitudes are responding to positive directional selection for increased corolla width over time. Corolla length displayed a non-significant negative relationship between *𝛿t* and the starting mean trait value in 2003 (*t*) (R = −0.46, p = 0.083) and no indication that this relationship significantly changed following the inclusion of latitude in the model.

**Figure 5.**
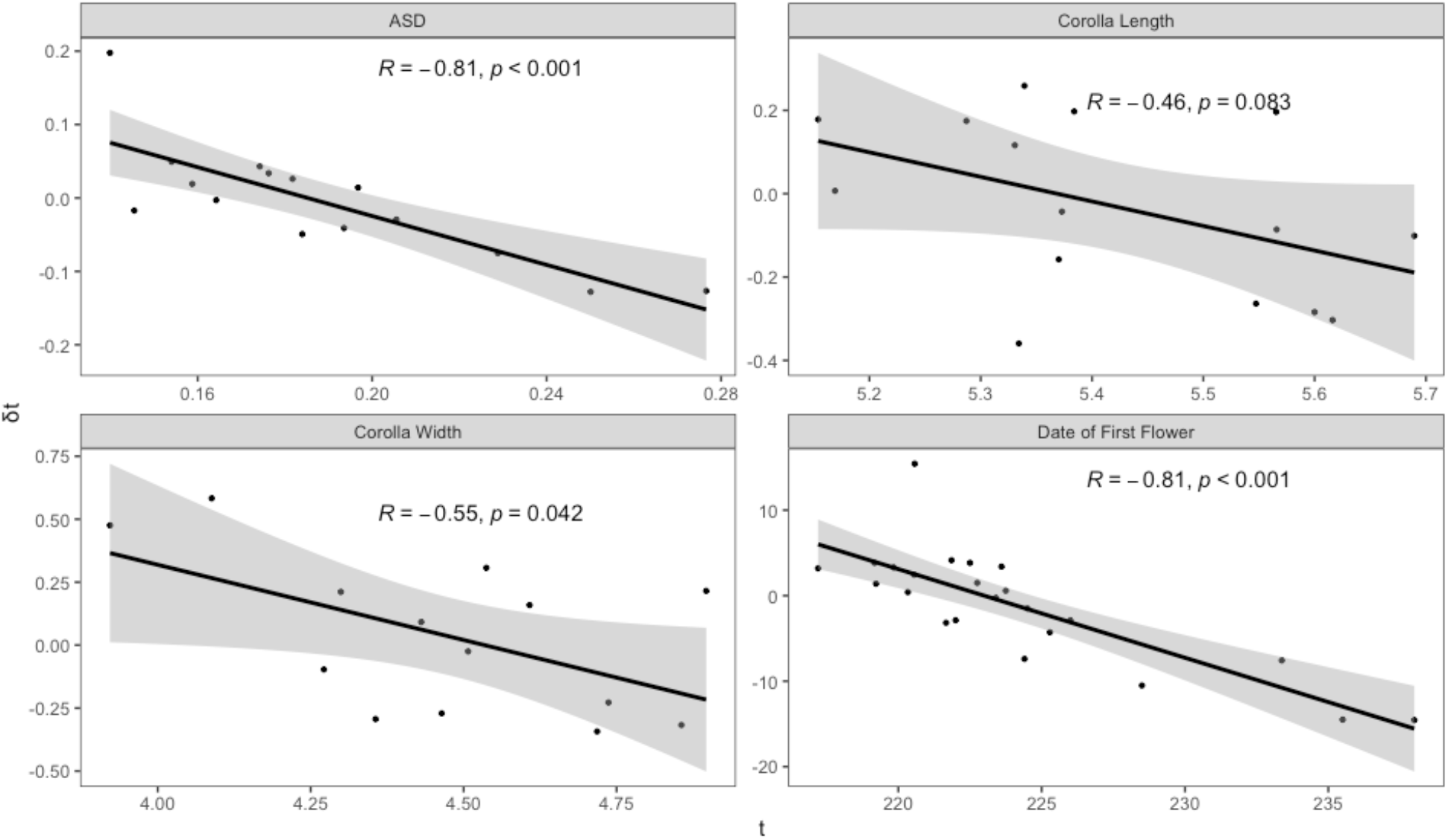
Linear regression showing the predictability of degree of change in trait value from 2003 to 2012 by the starting trait value in 2003. Each point in the regression is the mean trait value for a population, with date of first flower shown only for the first flowering wave. *t* represents the starting mean value in 2003, and 𝛿*t* shows the degree of change in mean value as the mean in 2003 subtracted from the mean in 2012.

Finally, both the anther-stigma distance and flowering time of the first flowering wave showed a highly significant and negatively sloped relationship between the change in the mean value of the trait and the starting mean value (ASD: R = −0.81, p = 2.0 × 10^−4^; first flowering wave: R = −0.81, p = 2.5 × 10-5). This relationship was significant regardless of whether latitude was included as an explanatory variable in an analysis of variance (ASD: F = 26.91, numDF = 1, denDF = 11, p = 3.004 × 10^−4^, first flowering wave: F = 38.31, numDF = 1, denDF = 19, p = 6.022 × 10^−6^). Neither trait showed evidence of reduced variation over time (Table S3). For ASD, *𝛿*t values are evenly distributed around 0 whereas *𝛿*t values for first flower are predominantly below zero (8 populations above, 15 populations below). This indicates that ASD is evolving under negative-frequency dependent selection whereas the day of first flowering (first flowering wave) is responding to positive directional selection (note that earlier flowering corresponds to a negative shift in trait value; Table S3, Figure 5).

While we did not examine the *𝛿*t/t relationship for floral reward traits due to low sample size (N = 4 populations), we note that the percent change in the phenotypic coefficient of variation (PCV) values show a significant decrease in both floral reward traits between 2003 to 2012 (°Brix: t = 1.970, p = 0.05; pollen number: t = 2.399, p = 0.01; Table S3), in alignment with the idea that these traits are responding to selection.

## DISCUSSION

Climate change encompasses both direct (abiotic) and indirect (biotic interactions) forces of selection, the effects of which can manifest in a range of growth and reproductive responses that maintain demographic performance despite substantial environmental change (Eckert *et al*., 2010). Uncertainty over climate response is especially acute for mixed-mating species, where, in the face of global pollinator declines and shifting suites of abiotic variables, both selection for increased outcrossing (Bishop *et al*., 2017) and selection for increased selfing (Jones *et al*., 2013) are possible adaptive responses. Our results show *I. purpurea* is evolving broader corollas with some evidence for increased floral rewards. We found no indication that anther-stigma distance decreased over time across examined populations, which would be expected if populations were evolving higher rates of selfing. Overall, our findings are aligned with the expectation of increased investment in pollinator attraction traits, especially at the northernmost populations, rather than increased rates of self-pollination.

Although patterns of trait change across each of the traits were compelling, corolla width showed the most dramatic overall increase in trait value from 2003 to 2012, as well as a decrease in the phenotypic coefficient of variation – together suggesting corolla width is responding to positive selection for increased size. An important nuance of this conclusion is that the evidence for such change in this trait was largely driven by the northernmost populations. Specifically, the use of a spatially explicit model for predicting the change in trait value over time(*𝛿t*) showed that *𝛿t* was better explained by the interaction between latitude and *t* rather than by *t* alone. Additionally, by removing the three southernmost populations in this regression, we found that the slope of *𝛿t/t* became positive (albeit p = 0.14), demonstrating again that corolla width changes are much larger in the north. Thus, the strongly significant spatial-temporal change in corolla width as well as decrease in variation for the trait are highly suggestive of directional selection occurring at northern latitudes. We found some indication of a temporal increase in corolla length, but believe this pattern is more likely due to the strong correlation between corolla width and length (r = 0.61 in 2003 and 0.59 in 2012, p < 2 × 10^−16^), rather than due to direct selection on corolla length. While corolla length plays an important role in pollination efficiency in some plant species with specialist pollinators (Naghiloo *et al*., 2021; Faure *et al*., 2022), *I. purpurea* is a generalist-pollinated plant, and the observed change in corolla length is so slight (increased 0.4 mm) that it is unlikely that this is a biologically significant effect or that pollinator efficiency is impacted by corolla length. Overall, our finding of a change in corolla width as a possible adaptive response to climate change aligns with previous evidence that corolla width is responsive to abiotic changes such as water availability and temperature, as well as changes in pollinator populations (Schueller, 2007; Campbell & Powers, 2015; Gallagher & Campbell, 2017). However, tracking floral traits over time remains rare, and, in contrast to our results, multiple other studies have suggested an increased investment in selfing in response to climate change and pollinator declines (Cheptou, 2019; Busch *et al*., 2022) with rarer instances of increased outcrossing (Bishop *et al*., 2017)

Our phenology and anther-stigma distance results are similar to responses across other species and previous work in *I. purpurea*, respectively. Phenology, measured here as the date of first flower, has repeatedly shown a shift to earlier flowering dates in a number of plant species (Bock *et al*., 2014; Moore & Lauenroth, 2017; Wolf *et al*., 2017; Büntgen *et al*., 2022), and this is also the case in *I. purpurea*. We found some evidence of directional selection toward earlier flowering within the first wave of flower emergence, particularly at northern latitudes. However, the bimodal distribution of first flowering dates demonstrates that earlier flowering is not captured fully within a single wave, rather, the mechanism underlying this shift is a greater proportion of individuals flowering in the first wave in 2012, rather than a shift of both peaks to earlier dates while retaining a bimodal nature. In fact, while all 23 populations have some individuals that flower in the first wave and some in the second in 2003, three of the populations flower entirely in the first wave in 2012. For anther-stigma distance, our results strongly point to negative frequency dependent selection acting on this trait over time, a result corroborating previous empirical work in a single experimental population which showed that outcrossing success in *I. purpurea* is negatively frequency dependent (Chang & Rausher, 1998). Our explicit spatial-temporal model for ASD showed no effect of latitude on ASD values, indicating that, unlike corolla width, selection is consistent across space.

While our data potentially indicate that both of the floral rewards – nectar sucrose content (°Brix) and pollen number – change over time (*i*.*e*., significant reduction in variation between years for both traits; mean trait increase for nectar sugar in northernmost populations), due to a low number of populations examined in this study (N = 4), we cannot assess selection on them. We likewise did not examine the potential that such changes are correlated to, and perhaps evolving along with corolla width again due to sample size limitations. Furthermore, both sucrose content and pollen count are only one component of potential rewards. In the case of nectar, volume and thus sugar concentration remain unaccounted for, while pollen count indicates little of pollen protein content. Nonetheless, it appears likely that there is a temporal increase in investment in pollinator attraction, and that this result is driven by populations at northern latitudes. Changes in floral rewards in response to climate change also align with previous findings indicating that both temperature and water availability can alter nectar volume and sugar content (Descamps *et al*., 2018, 2021; Phillips *et al*., 2018) as well as pollen count and viability (Bishop *et al*., 2017; Descamps *et al*., 2021).

This is the first paper to use the resurrection approach to examine the potential that traits responsible for plant-pollinator interactions may be evolving over time, concomitant to decreases in pollinator abundance and dramatic environmental changes due to a changing climate. While a unique feature of the resurrection approach is that it allows for comparisons of populations exposed to the multifactorial suite of selective pressures associated with climate change in the field (Thomann *et al*., 2013), the resurrection approach typically does not identify the causative agent(s) of selection, meaning that we will need to perform future direct manipulations of abiotic and biotic factors to determine which agents of selection are acting on corolla width and other floral traits. However, with some notable exceptions (Inouye, 2008; Franks, 2011; Anderson et al., 2012; Thomann et al., 2015; Rauschkolb et al., 2022), relatively few studies investigating adaptation to climate change capture adaptive responses from field settings, showcasing the power of the approach we have taken here. Another important caveat to the resurrection approach is bias introduced by the “invisible fraction” – that is, when nonrandom mortality of stored seeds creates bias in measurements of phenotypic traits due to association between traits of interest and traits related to germination success (Weis, 2018). In this study, germination rates between 2003 and 2012 were very high and slightly higher in the older seeds (87% in 2003, 84% in 2012), such that germination failure is unlikely to be related to seed traits affecting storage survival and bias in trait measurements is expected to be trivial (Weis, 2018). Despite these caveats, our results are compelling in that they are in alignment with phenological shifts in other plant species ((Byers, 2017; Renner & Zohner, 2018; Gérard *et al*., 2020; Soares *et al*., 2021)) and a broad range of work showing northern populations tend to show more dramatic evolutionary responses to climate change (Parmesan, 2007; Bonebrake *et al*., 2010; Post *et al*., 2018)

In summary, we show that, in addition to well-documented shifts to earlier flowering phenology, floral architecture and rewards can also play significant roles in the evolutionary response to contemporary environmental change. Populations of *I. purpurea* distributed across the southeast United States demonstrate a significant temporal increase in corolla size as well as potential for increased investment in floral rewards, all of which are driven primarily by populations at more northern latitudes. In addition, we show that the integration of phenotypic trait changes over time, measurement of variation, and spatial modeling can be used to detect signals of selection on phenotypic traits, notably, the presence of balancing selection on anther-stigma distance, and a probable instance of spatially divergent directional selection on floral architecture.

## ACKNOWLEDGMENTS

The authors thank Adam Kuester, Malia Santos and Laurel Wellman for caring for greenhouse plants and taking measurements of floral morphology, nectar sucrose, and pollen counts, respectively. The authors have no conflict of interest to declare.

## AUTHOR CONTRIBUTIONS

RSB and SB conceived and designed the study with RSB acting as the advisor. SMC collected phenology data, and SB analyzed the data from all three greenhouse experiments. SB and RSB drafted the initial version of the manuscript and all authors contributed to later versions of the manuscript.

