## Supplemental File 1 (Supplemental Tables 1, 2, 3, and 4) for "Not just flowering time: A resurrection approach shows floral attraction traits are changing over time"

| | | | | | | Number of Plants | | | Number of Flowers | | Number of Maternal Lines |
|---|---|---|---|---|---|---|---|---|---|---|---|
| **ID** | **Population Name** | **Year** | **State** | **Latitude** | **Longitude** | **Morphology** | **Phenology** | **Rewards** | **Morphology** | **Rewards** | **Rewards** |
| 1 | billings | 2003 | TN | 35.77524 | -85.90342 | 10 | NA | NA | 55 | NA | NA |
| 1 | billings | 2012 | TN | 35.77524 | -85.90342 | 22 | NA | NA | 191 | NA | NA |
| 2 | bergaw1 | 2003 | NC | 34.59571 | -77.92748 | 13 | 14 | NA | 68 | NA | NA |
| 2 | bergaw1 | 2012 | NC | 34.59571 | -77.92748 | 15 | 2 | NA | 84 | NA | NA |
| 4 | chicken road | 2003 | NC | 34.55667 | -79.12560 | 16 | 12 | NA | 74 | NA | NA |
| 4 | chicken road | 2012 | NC | 34.55667 | -79.12560 | 17 | 12 | NA | 60 | NA | NA |
| 5 | clarendon1 | 2003 | SC | 33.85988 | -79.90907 | NA | 13 | NA | NA | NA | NA |
| 5 | clarendon1 | 2012 | SC | 33.85988 | -79.90907 | NA | 1 | NA | NA | NA | NA |
| 8 | darlington2 | 2003 | SC | 34.29720 | -79.99126 | 16 | 10 | NA | 90 | NA | NA |
| 8 | darlington2 | 2012 | SC | 34.29720 | -79.99126 | 12 | 11 | NA | 46 | NA | NA |
| 9 | duplin east | 2003 | NC | 34.92404 | -77.79617 | NA | 13 | 26 | NA | 179 | 8 |
| 9 | duplin east | 2012 | NC | 34.92404 | -77.79617 | NA | 4 | 8 | NA | 65 | 6 |
| 10 | duplin west | 2003 | NC | 34.98316 | -78.03931 | NA | 9 | NA | NA | NA | NA |
| 10 | duplin west | 2012 | NC | 34.98316 | -78.03931 | NA | 2 | NA | NA | NA | NA |
| 11 | grimsley | 2003 | NC | 34.52714 | -78.75670 | NA | 5 | NA | NA | NA | NA |
| 11 | grimsley | 2012 | NC | 34.52714 | -78.75670 | NA | 16 | NA | NA | NA | NA |

| | | | | | | | | | | | |
|---|---|---|---|---|---|---|---|---|---|---|---|
| 12 | florence | 2003 | SC | 34.14581 | -79.86531 | 18 | 12 | 28 | 78 | 195 | 8 |
| 12 | florence | 2012 | SC | 34.14581 | -79.86531 | 13 | 11 | 31 | 83 | 222 | 8 |
| 14 | hare road | 2003 | NC | 35.42476 | -77.91712 | 16 | 14 | 32 | 106 | 208 | 8 |
| 14 | hare road | 2012 | NC | 35.42476 | -77.91712 | 17 | 11 | 31 | 97 | 213 | 8 |
| 15 | horry1 | 2003 | SC | 34.10421 | -79.07373 | 15 | 10 | NA | 94 | NA | NA |
| 15 | horry1 | 2012 | SC | 34.10421 | -79.07373 | 12 | 13 | NA | 53 | NA | NA |
| 16 | horry2 | 2003 | SC | 34.10535 | -79.18323 | 13 | 9 | NA | 101 | NA | NA |
| 16 | horry2 | 2012 | SC | 34.10535 | -79.18323 | 29 | 11 | NA | 177 | NA | NA |
| 17 | marion1 | 2003 | SC | 34.15915 | -79.27291 | 15 | 11 | NA | 97 | NA | NA |
| 17 | marion1 | 2012 | SC | 34.15915 | -79.27291 | 18 | 12 | NA | 96 | NA | NA |
| 18 | marion2 | 2003 | SC | 34.15659 | -79.27027 | NA | 11 | NA | NA | NA | NA |
| 18 | marion2 | 2012 | SC | 34.15659 | -79.27027 | NA | 13 | NA | NA | NA | NA |
| 19 | mckinnon | 2003 | NC | 34.50819 | -78.70899 | 17 | 13 | NA | 135 | NA | NA |
| 19 | mckinnon | 2012 | NC | 34.50819 | -78.70899 | 20 | 8 | NA | 195 | NA | NA |
| 21 | new hope | 2003 | NC | 35.36982 | -77.87731 | NA | 13 | NA | NA | NA | NA |
| 21 | new hope | 2012 | NC | 35.36982 | -77.87731 | NA | 11 | NA | NA | NA | NA |
| 22 | oldkenley | 2003 | NC | 36.14360 | -78.05342 | 18 | 12 | NA | 125 | NA | NA |
| 22 | oldkenley | 2012 | NC | 36.14360 | -78.05342 | 27 | 10 | NA | 279 | NA | NA |
| 23 | snakes | 2003 | TN | 35.06791 | -86.62955 | NA | 3 | NA | NA | NA | NA |
| 23 | snakes | 2012 | TN | 35.06791 | -86.62955 | NA | 11 | NA | NA | NA | NA |
| 25 | starlight | 2003 | NC | 34.61636 | -79.05167 | 17 | 12 | NA | 89 | NA | NA |
| 25 | starlight | 2012 | NC | 34.61636 | -79.05167 | 1 | 5 | NA | 4 | NA | NA |

| | | | | | | | | | | | |
|---|---|---|---|---|---|---|---|---|---|---|---|
| 26 | spears soy | 2003 | TN | 35.53341 | -85.95190 | *NA* | 3 | *NA* | *NA* | *NA* | *NA* |
| 26 | spears soy | 2012 | TN | 35.53341 | -85.95190 | *NA* | 9 | *NA* | *NA* | *NA* | *NA* |
| 28 | sumter2 | 2003 | SC | 34.09792 | -80.37771 | 16 | 9 | 25 | 62 | 187 | 8 |
| 28 | sumter2 | 2012 | SC | 34.09792 | -80.37771 | 13 | 9 | 29 | 60 | 199 | 8 |
| 29 | tarheal | 2003 | NC | 34.70513 | -78.73890 | 14 | 12 | *NA* | 63 | *NA* | *NA* |
| 29 | tarheal | 2012 | NC | 34.70513 | -78.73890 | 10 | 9 | *NA* | 65 | *NA* | *NA* |
| 30 | willis corn | 2003 | TN | 35.31105 | -85.94500 | *NA* | 11 | *NA* | *NA* | *NA* | *NA* |
| 30 | willis corn | 2012 | TN | 35.31105 | -85.94500 | *NA* | 9 | *NA* | *NA* | *NA* | *NA* |
| 31 | vervilla | 2003 | TN | 35.60848 | -85.84638 | *NA* | 10 | *NA* | *NA* | *NA* | *NA* |
| 31 | vervilla | 2012 | TN | 35.60848 | -85.84638 | *NA* | 10 | *NA* | *NA* | *NA* | *NA* |
| 32 | walnut grove | 2003 | TN | 35.09936 | -86.22551 | 6 | *NA* | *NA* | 57 | *NA* | *NA* |
| 32 | walnut grove | 2012 | TN | 35.09936 | -86.22551 | 10 | *NA* | *NA* | 52 | *NA* | *NA* |

**Supplementary Table S1.** Sampling localities included in the phenotypic analyses including geographic coordinates, the number of plants included for each of the three greenhouse experiments, the number of flowers measured for the floral morphology and floral rewards experiments, and the number of maternal lines used in the floral rewards experiment. Abbreviations: TN: Tennesses, NC: North Carolina, SC: South Carolina

| | | Corolla Width | Corolla Length | ASD | First Flowering Wave | Second Flowering Wave | °Brix | Pollen Count |
|---|---|---|---|---|---|---|---|---|
| **Trait ~ Year*Latitude + (1\|Population)** | | | | | | | | |
| **Year** | numDF | 1 | 1 | 1 | 1 | 1 | 1 | 1 |
| | denDF | 12.10 | 11.77 | 9.67 | 300.97 | 139.97 | 1.94 | 1.90 |
| | F | 7.093 | 10.472 | 4.42 | 3.300 | 0.232 | 0.003 | 0.028 |
| | p | 0.0205** | 0.007** | 0.659 | 0.070* | 0.631 | 0.961 | 0.883 |
| **Latitude** | numDF | 1 | 1 | 1 | 1 | 1 | 1 | 1 |
| | denDF | 2662.85 | 2781.94 | 2468.51 | 22.151 | 22.12 | 200.63 | 163.41 |
| | F | 16.850 | 0.041 | 1.587 | 1.169 | 6.194 | 1.94 | 2.187 |
| | p | 4.167e-05** | 0.840 | 0.072 | 0.291 | 0.021** | 0.016** | 0.141 |
| **Year*Latitude** | numDF | 1 | 1 | 1 | 1 | 1 | 1 | 1 |
| | denDF | 519.82 | 923.79 | 1576.65 | 300.97 | 139.94 | 60.45 | 33.25 |
| | F | 23.388 | 0.580 | 2.633 | 3.339 | 0.225 | 4.59 | 2.180 |
| | p | 1.747e-06** | 0.447 | 0.641 | 0.069* | 0.636 | 0.036** | 0.149 |
| **$\delta t$ ~ t*Latitude** | | | | | | | | |
| **t** | numDF | 1 | 1 | 1 | 1 | 1 | NA | NA |
| | denDF | 11 | 11 | 11 | 19 | 16 | NA | NA |

| | | | | | | | | |
|---|---|---|---|---|---|---|---|---|
| | F | 2.600 | 3.507 | 26.905 | 38.307 | 6.291 | NA | NA |
| | p | 0.14 | 0.088* | 3.004e-4** | 6.022e-6** | 0.023** | NA | NA |
| **Latitude** | numDF | 1 | 1 | 1 | 1 | 1 | NA | NA |
| | denDF | 11 | 11 | 11 | 19 | 16 | NA | NA |
| | F | 2.654 | 1.277 | 3.055 | 0.789 | 1.157 | NA | NA |
| | p | 0.13 | 0.283 | 0.108 | 0.386 | 0.298 | NA | NA |
| **t*Latitude** | numDF | 1 | 1 | 1 | 1 | 1 | NA | NA |
| | denDF | 11 | 11 | 11 | 19 | 16 | NA | NA |
| | F | 6.058 | 0.671 | 0.345 | 3e-4 | 0.792 | NA | NA |
| | p | 0.032** | 0.430 | 0.569 | 0.987 | 0.387 | NA | NA |

**Table S2.** Linear mixed models showing the influence of latitude on temporal changes in trait value and the predictability of $\delta t$ by $t$. The first model shows significance of latitude, sampling year, and the interaction of the two on all trait values, reported as a truncated p-value based on Satterwaite's degrees of freedom method for a type III ANOVA. For floral morphology and phenology, latitude is included as a fixed effect while population is used as a random effect to control for longitude. The model for floral rewards additionally uses maternal line as a nested random effect within population (Trait ~ Year*Latitude + (1|Pop/ML)). The second model shows the interactive effect of latitude and starting population mean trait value on the degree of change in mean trait value for a population. Significance ($p<0.05$) is marked with a double asterisk** and relationships trending significant ($0.05 < p < 0.1$) are marked with a single asterisk*.

| | Year | Range | Mean | SD | PCV (%) | % Change PCV | Significance (p) |
|---|---|---|---|---|---|---|---|
| **Corolla Width** | 2003 | 1.8-7.5 | 4.503 | 0.993 | 21.976 | NA | NA |
| | 2012 | 1-7.1 | 4.764 | 0.993 | 20.849 | 5.406 | 0.063 |
| **Corolla Length** | 2003 | 2.8-8 | 5.431 | 0.616 | 11.463 | NA | NA |
| | 2012 | 2.2-7 | 5.474 | 0.6165 | 11.259 | 1.812 | 0.642 |
| **ASD** | 2003 | -2.7-1.3 | 0.116 | 0.256 | 212.296 | NA | NA |
| | 2012 | -1-1.15 | 0.108 | 0.201 | 186.923 | 13.57 | 0.387 |
| **Flowering Wave 1** | 2003 | 213-240 | 223.2 | 8.112 | 3.629 | NA | NA |
| | 2012 | 214-241 | 223 | 7.466 | 3.343 | 8.555 | 0.211 |
| **Flowering Wave 2** | 2003 | 256-305 | 277.8 | 13.82 | 4.959 | NA | NA |
| | 2012 | 257-307 | 278.3 | 14.26 | 5.103 | -2.822 | 0.753 |
| **Nectar Sucrose Content** | 2003 | 0-21 | 6.524 | 3.056 | 46.838 | NA | NA |
| | 2012 | 0-22 | 6.805 | 2.903 | 42.66 | 9.794 | 0.048 |
| | 2003 | 3-508 | 191.5 | 59.25 | 30.925 | NA | NA |

| | | | | | | | |
|---|---|---|---|---|---|---|---|
| **Pollen Count** | 2012 | 2-493 | 200.8 | 49.92 | 24.855 | 24.42 | 0.016 |

**Table S3.** Range, least squares mean, and standard deviation of floral trait values are shown for each year. Floral morphology traits are measured in centimeters, flowering dates use a Julian calendar, nectar sucrose content is measured as °Brix, and pollen count is a total number of pollen grains found on a single anther. Also shown is the phenotypic coefficient of variation (PCV) determined using a bootstrapped resampling procedure with 10000 draws, percent change of PCV from 2003 to 2012, and the p-value associated with a two-sided independent t-test comparing levels of phenotypic variation in 2003 to 2012 for each trait.

| | CW:CL | | CW:ASD | | CL:ASD | | CW:FF | | CL:FF | | ASD:FF | |
|---|---|---|---|---|---|---|---|---|---|---|---|---|
| | 2003 | 2012 | 2003 | 2012 | 2003 | 2012 | 2003 | 2012 | 2003 | 2012 | 2003 | 2012 |
| Correlation Coefficient (r) | 0.611 | 0.596 | 0.035 | -0.030 | -0.055 | -0.129 | -0.237 | -0.008 | -0.188 | 0.081 | 0.364 | -0.174 |
| Significance (p) | <2.2e-16 | <2.2e-16 | 0.223 | 0.253 | 0.047 | 3.681e-07 | 0.245 | 0.970 | 0.359 | 0.695 | 0.068 | 0.394 |

**Table S4.** Pearson's correlation coefficients and associated p-values for floral morphology traits and phenology. Brix and pollen count excluded due to low population sampling. Since floral traits and phenology are not paired data, correlation is calculated with population means, otherwise correlation coefficients and p-values are calculated from raw measurements. Repeating a Pearson's correlation test on floral traits using population means yields a higher correlation between floral traits. Abbreviations: CW = Corolla Width, CL = Corolla Length, ASD = Anther Stigma Distance, FF = date of first flower
